# Noise-resistant developmental reproducibility in vertebrate somite formation

**DOI:** 10.1101/247783

**Authors:** Honda Naoki, Ryutaro Akiyama, Shin Ishii, Yasumasa Bessho, Takaaki Matsui

## Abstract

The reproducibility of embryonic development is remarkable, although molecular processes are intrinsically stochastic at the single-cell level. How the multi-cellular system resists the inevitable noise to acquire developmental reproducibility constitutes a fundamental question in developmental biology. Toward this end, we focused on vertebrate somitogenesis as a representative system, because somites are repeatedly reproduced within a single embryo whereas such reproducibility is lost in segmentation clock gene-deficient embryos. However, the effect of noise on developmental reproducibility has not been fully investigated, because of the technical difficulty in manipulating the noise intensity in experiments. In this study, we developed a computational model of ERK-mediated somitogenesis, in which bistable ERK activity is regulated by an FGF gradient, cell-cell communication, and the segmentation clock, subject to the intrinsic noise. The model simulation generated our previous *in vivo* observation that the ERK activity was distributed in a step-like gradient in the presomitic mesoderm, and its boundary was posteriorly shifted by the clock in a stepwise manner, leading to regular somite formation. Here we showed that this somite regularity was robustly maintained against the noise. Removing the clock from the model predicted that the stepwise shift of the ERK activity occurs at irregular timing with irregular distance owing to the noise, resulting in somite size variation. Through theoretical analysis, we presented a mechanism by which the clock reduces the inherent somite irregularity observed in clock-deficient embryos. Therefore, this study indicates a novel role of the segmentation clock in noise-resistant developmental reproducibility.

## Introduction

Embryonic development in multicellular organisms is highly reproducible. Conversely, biochemical reactions at the single-cell level are intrinsically stochastic owing to the low copy numbers of expressed molecules (1–3). Thus, it is believed that organisms have a homeostatic mechanism that can resist the stochasticity of cells, while maintaining the precision and reproducibility required for normal development (4). However, how the noise affects development *in vivo* remains unclear, because of the technical difficulty in manipulating the noise intensity in experiments. We approached this issue through computational modeling, especially focusing on vertebrate somitogenesis as a model biological system. Equal-sized somites are continuously produced within a single embryo (Fig. 1*A* and *B*); thus, reproducibility can be easily evaluated in vertebrate somitogenesis within single, rather than among different, embryos.

**Fig. 1.**
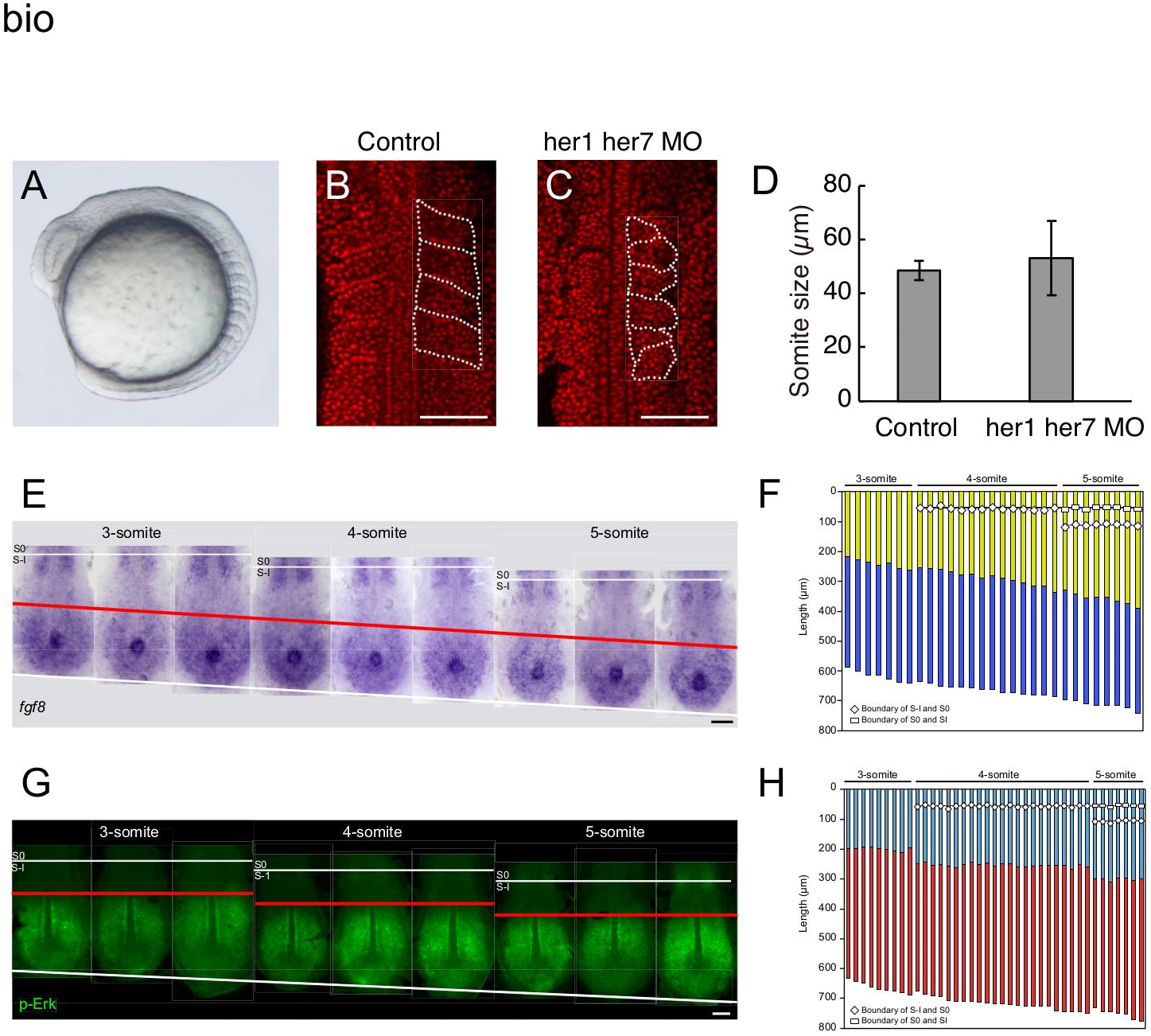
Somite reproducibility and stepwise ERK activity regression for somite formation. *(A)* Lateral view of a wild-type zebrafish embryo during somitogenesis. (*B, C*) Somite morphology in wild-type (B) or clock-deficient embryos co-injected with *her1*- and *her7*-morpholinos. Nuclei are stained by propidium iodide (red). Somites are outlined by dotted lines. Scale bar: 100 µm. (*D*) Somite size variation in control and clock-deficient embryos (n=16 each). Data represent the means and standard deviations. (*E*) Representative dorsal view of PSM *fgf8a* mRNA expression at 3- to 5-somite stages. Red line, *fgf8a* gradient boundary. Scale bars: 100 µm. (*F*) Quantitative presentation of *fgf8a* expression. Embryos (n = 29) were arranged in order of developmental stages, as estimated by somite number and PSM length. High (purple stripe) and low (yellow stripe) *fgf8a* expression domains in each embryo. (G) Representative dorsal view of PSM p-ERK distribution at 3- to 5-somite stages. Red line, p-ERK boundary. (*H*) Quantitative presentation of p-Erk distribution. Embryos (n = 39) were arranged as in (*F*). ON (red stripe) and OFF (blue stripe) Erk activity regions in each embryo. Panels *B*, *C*, and *E-H* are modified from our original paper (18).

Somites are generated by periodic segmentation of the anterior end of the presomitic mesoderm (PSM), which serves as the foundation for the metameric structures of the vertebrate body (5–7). The most important concept of somitogenesis is the clock and wavefront (CW) model proposed in the 1970s (8). This model comprises two signals; namely, the oscillatory clock and posteriorly traveling wavefront in the growing PSM. It predicts that, when the wavefront meets the clock signal, PSM cells start to differentiate into somite cells, leading to somite individualization. After extensive investigation, the molecular bases of CW were identified in various vertebrates (6, 9). Clock genes show oscillatory expression in the PSM: *Hes7* (a *Hes* family transcription factor) and *Lunatic fringe* (*Lfng*, a Notch effector) in the mouse (10, 11), and *her1* and *her7* (*Hes* family transcription factors) and *deltaC* (a Notch ligand) in the zebrafish (12, 13). Furthermore, morphogen gradients in the PSM function as the wavefront; fibroblast growth factor (FGF) and Wnt signaling produce gradients that weaken at the anterior PSM and continuously move toward the posterior end alongside tail elongation (6, 14–17).

However, the effect of innate noise on somitogenesis developmental reproducibility has been outside of the scope of the CW model. Thus, how anti-noise robustness can be acquired remains as an important question for both improving the CW model and elucidating a strategy of noise-resistant developmental reproducibility. Loss-of-function phenotypes of clock genes may provide clues for this issue; clock-deficient mouse (*Hes7*-deficient) and zebrafish (*her1* and *her7* double morphants and/or mutants) embryos still form somites, albeit with variable size (Fig. 1*C* and *D*), resulting in severe skeletal and segmental defects (10, 18, 19). This suggest that the clock somehow controls somite size regularity by reducing the noise effect, and that noise may induce variably-sized somite formation in a clock-independent manner. Currently, it remains unclear how irregular somites form in clock-deficient embryos and how their irregularity is suppressed in control embryos.

To understand how segmental pre-patterns are determined, we previously investigated FGF/ERK signaling by collecting stained zebrafish embryos (18). We found that the gentle *fgf8a* gradient (wavefront) was converted into a step-like gradient of ERK activity with a sharp border, which was regularly shifted by the segmentation clock in a stepwise manner (Fig. 1*E*-*H*). This suggested that ERK integrates clock and wavefront signals and establishes a stepwise pattern within the uniform PSM, which is required for somite individualization. However, an ERK activity stepwise pattern could not be detected in clock-deficient embryos (*her1* and *her7* double morphants), as the spatiotemporal resolution was quite low in our static analyses.

The aim of this study is to clarify how developmental reproducibility is acquired against inevitable noise in somitogenesis. By developing a computational model of ERK-mediated somitogenesis, we investigated the effect of the noise during somitogenesis. This model generated the stepwise ERK activity shift as observed in the PSM, which was robust against the noise. It also predicted that the shift occurred even in the absence of the clock, which could not be observed in our previous static analyses, although the timing and distance were irregular owing to the noise, resulting in reduced somite reproducibility. In summary, this study uncovered the clock-independent mechanism of irregular somite formation and also proposed a novel concept for a clock-dependent mechanism of noise-resistant developmental reproducibility in somite formation.

## Results

### Model of ERK-Mediated Somitogenesis

We first developed a computational model of ERK-mediated somitogenesis based on previous experimental findings (Fig. 2*A*-*C*; see Methods for details). The model described the ERK activity in the growing PSM. The ERK was regulated by several factors: the FGF gradient, positive feedback, cell-cell interactions, and the clock (Fig. 2*A*).

**Fig. 2.**
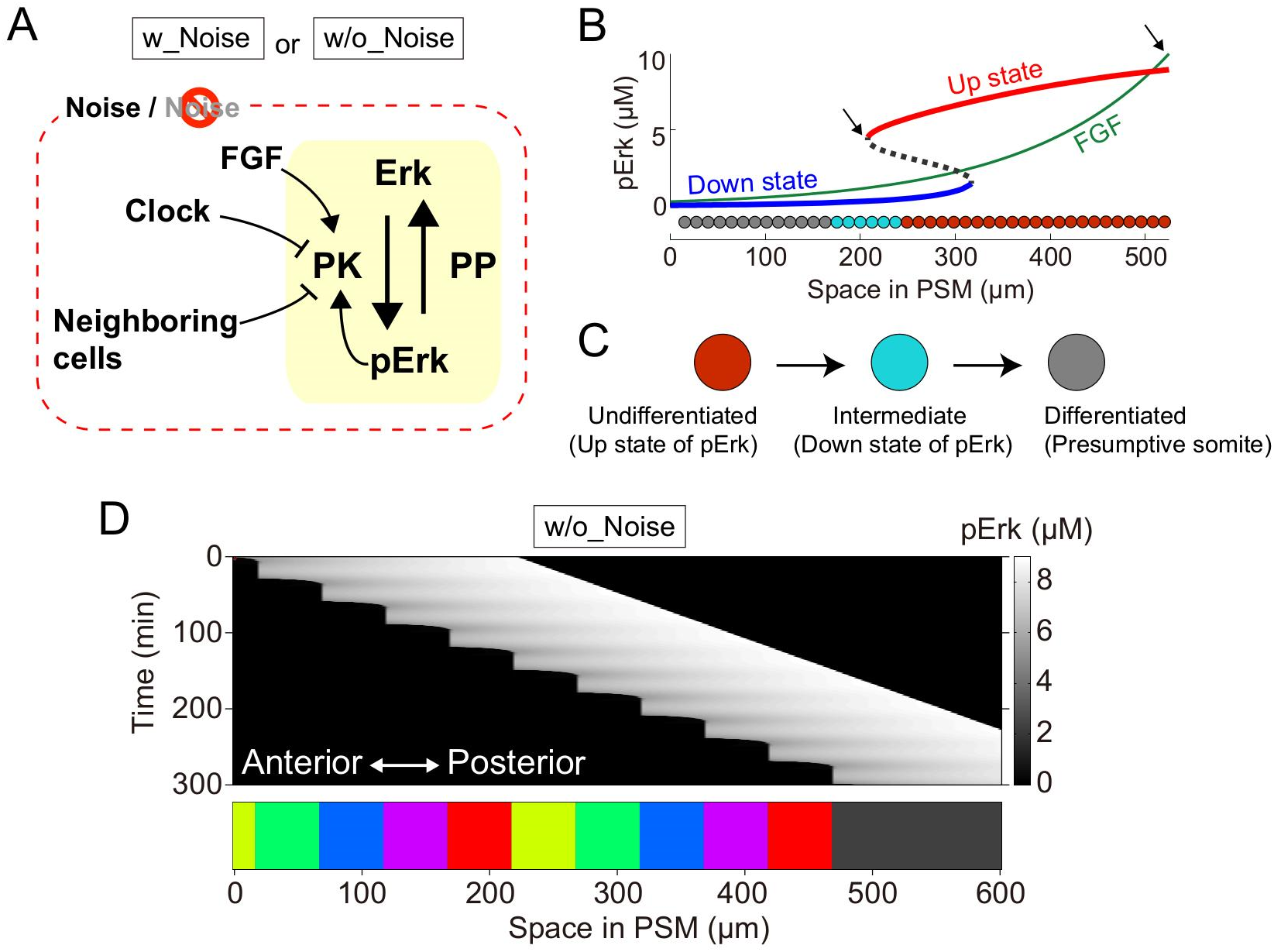
Model of ERK-mediated somite formation. (*A*) Schematic representation of ERK signaling. Our model includes the FGF gradient, positive feedback, segmentation clock, and cell-cell interaction. Simulations were performed with or without reaction noise. (*B*) PSM ERK activity bifurcation diagram. Up (red line) and down (blue line) ERK activity states. Black dashed line, unstable point between the up and down states; green line, FGF gradient. Because newly formed PSM cells are exposed to high FGF concentration, ERK activity is in the up state (right arrow). When cells reach the bifurcation point (left arrow), ERK activity rapidly shifts to the down state. (*C*) Cells are initially undifferentiated when ERK activity is in the up state. ERK activity decreases upon up-down transition, promoting cell entry to the intermediate state. The Intermediate cell cohort collectively differentiates into presumptive somites, if the up-down ERK activity state transition does not occur within a specific time interval. (*D*) Simulation result of ERK activity (upper panel) and the resultant somites (lower panel).

In the model, FGF is distributed in the PSM and activates ERK signaling (Fig. 2*B*). The FGF gradient continuously moves toward the posterior end alongside tail elongation, as observed in the PSM (Fig. 1*E* and *F*). As ERK activity exhibits a step-like distribution in the PSM with a sharp border (Fig. 1*G* and *H*), we assumed that ERK signaling positive feedback generates this bistability with the activity level switching between “up” (activated) and “down” (inactivated) states (Fig. 2*A* and *B*). Because somites can be spontaneously formed in a clock-independent manner via cell-cell interactions (20) and the ERK activity levels in a PSM cell and in adjacent PSM cells are similar (Fig. 1*G*), we assumed that ERK activity in neighboring cells is coupled via cell-cell interactions (Fig. 2*A*). In addition, we incorporated the effect of periodic ERK activity inhibition through the clock (Fig. 2*A*).

Based on the ERK dynamics, the PSM cells sequentially experienced three distinct states: undifferentiated, intermediate, and differentiated (committed as presumptive somites) (Fig. 2*C*). Undifferentiated PSM cells were characterized by an “up” ERK activity state (Fig. 2*B*, *red cells*). Upon ERK inactivation, the cells in a “down” state entered an intermediate state (Fig. 2B, *cyan cells*). If the up-down ERK activity state transition did not occur within a specific time interval, the intermediate cell cohort differentiated into presumptive somites (Fig. 2B, *grey cells*).

### Stepwise Shift of ERK Activity Distribution

Using this computational model, we simulated dynamic changes in ERK activity and ERK-mediated somitogenesis (Fig. 2*D*). A sharp ERK activity border was generated at a specific position in the PSM (Fig. 2*D*, *upper panel*) depending on the ERK activity bifurcation diagram (Fig. 2*B*). In addition, because of the continuous FGF gradient movement and the periodic clock effect on ERK activity, the border regressed in a stepwise manner, during which a group of cells collectively showed an up-down ERK activity transition (Fig. 2*D*, *upper panel*). Consequently, constant-sized somites were repeatedly formed (50.0 µm) (Fig. 2*D*, *lower panel*). Thus, our model successfully demonstrated stepwise ERK activity regression and normal somitogenesis, as observed *in vivo* (Fig. 1*A* and *B*).

### Robustness Against Noise

To explore the effect of intrinsic noise on ERK-mediated somitogenesis, we perturbed this system by adding the noise into the model (Fig. 3*A*). Even in the presence of the noise, the ERK activity border was generated and moved toward the posterior end in a stepwise manner, resulting in normal somitogenesis (50.0 ± 2.7 µm; C.V. = 0.05) (Fig. 3*B*). The system tolerated the noise upon further increase of the noise variance (*w*) over a wide range, (at least, from ×1 to ×5), leading to stepwise ERK activity regression and normal somite formation (Fig. 3*C*-*F*). These results therefore indicated that our model recapitulated normal somitogenesis in the presence of intrinsic noise, and that this system resisted and was robust against the noise.

**Fig. 3.**
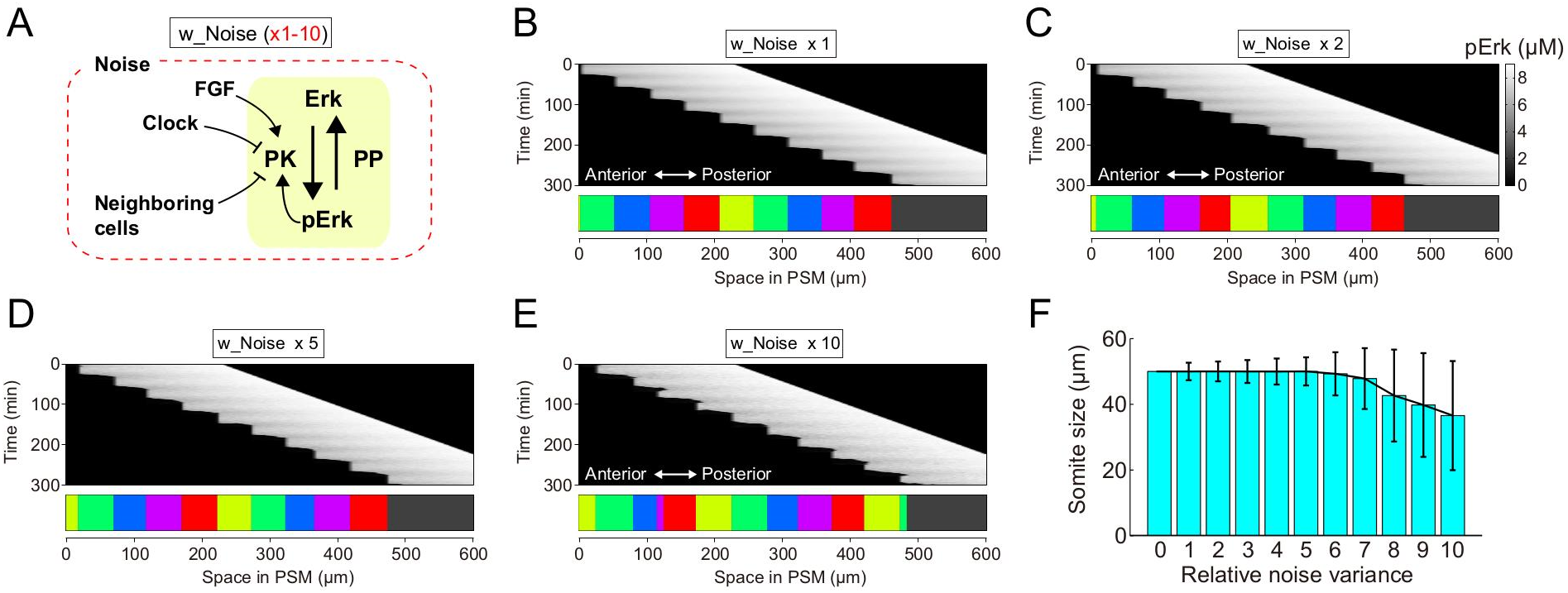
Somite size robustness against reaction noise. (*A*) Simulation settings. Simulations were performed with varying noise variance (i.e., *w* = 1–10 in Eq. 2). (*B-E*) Representative simulation results of Erk activity (upper panel) and the resultant somites (lower panel) upon noise variance at standard conditions (*B*) or increased 2 (*C*), 5 (*D*) or 10 (*E*) fold. (*F*) Noise effect on somite size. Formations of 200 somites were simulated by changing the noise variance from 0 to 10 fold. Data represent the means and standard deviations.

### Role of Bistability

To investigate how ERK-mediated somitogenesis obtains robustness against noise, we removed each component from our model. Upon removal of the positive feedback regulation of ERK (Fig. 4*A*), ERK activity showed a mild gradient within the PSM rather than a sharp border owing to the lack of bistability (Fig. 4*B*). In the absence of noise, the model, which lacks the positive feedback regulation, showed stepwise ERK activity regression and normal somite formation (Fig. 4*C*). However, in the presence of noise, stepwise ERK activity regression was retained, but somitogenesis became abnormal with co-existence of various-sized somites (Fig. 4*D*). These results suggest that the bistable nature of ERK dynamics, resulting from the positive feedback regulation, is required for robust somitogenesis in the presence of noise.

**Fig. 4.**
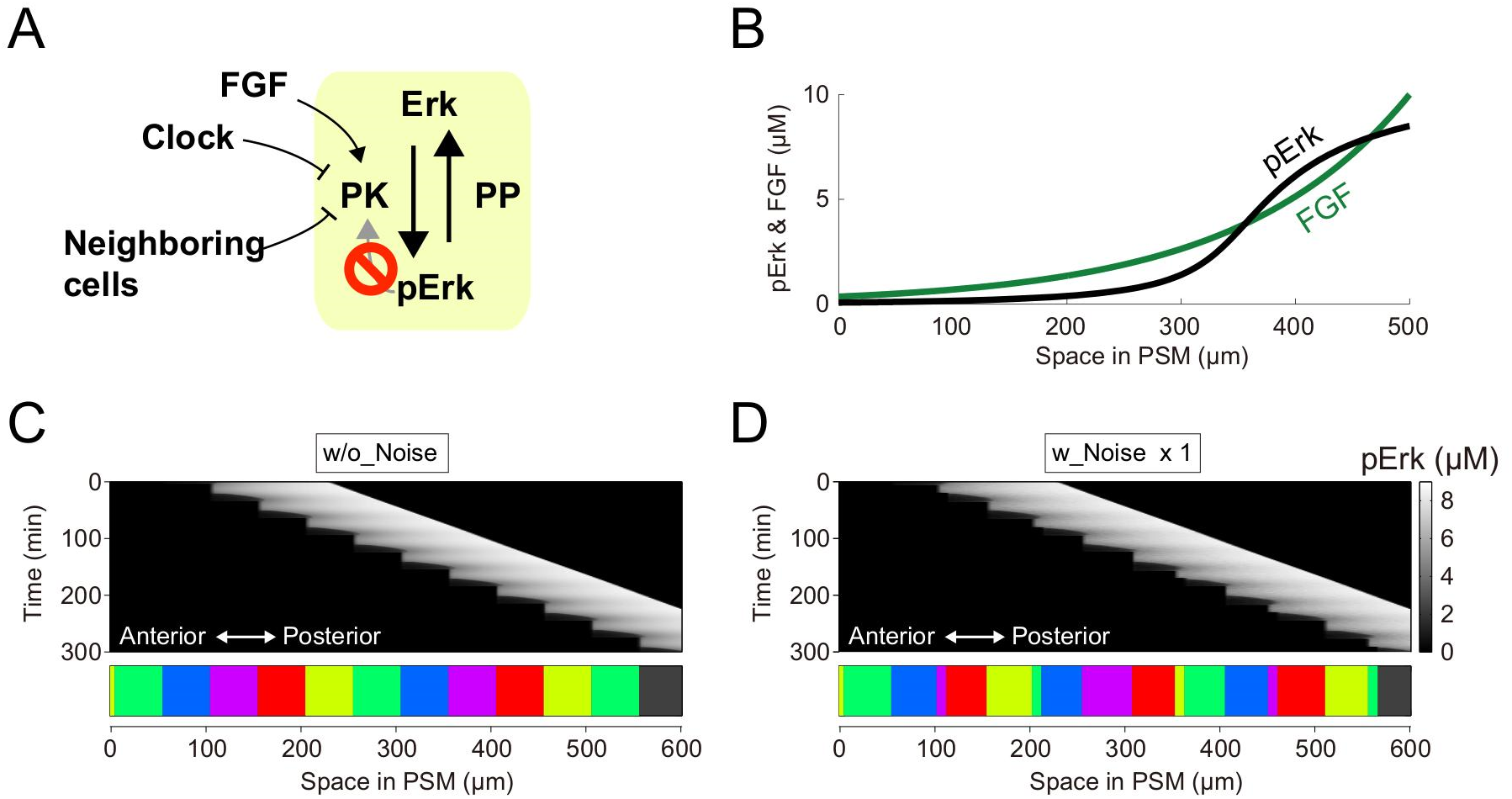
Removing Erk signaling positive feedback erases robustness against the noise. (*A*) Simulation settings. Simulations were performed upon positive feedback removal (i.e., *b*=0 in Eq. 3). (*B*) Without positive feedback, PSM Erk activity was monotonically distributed (black line). Green line, FGF gradient. (*C, D*) Schematic representation of simulation conditions (upper panels). Representative simulation results of Erk activity (middle panels) and the resultant somites (lower panels). Somites became regular (*C*) and irregular (*D*) in the absence and presence of noise, respectively, suggesting that positive feedback-induced bistability is necessary for somite reproducibility subject to noise.

### Role of Cell-Cell Interaction

To test the effects of cell-cell interactions on ERK-mediated somitogenesis and its robustness, we removed these from our model (Fig. S1*A*). In the absence of noise and cell-cell interactions, our model could generate stepwise ERK activity regression and normal somite formation (Fig. S1*B*). However, in the presence of noise, inhomogeneity of ERK activity among the PSM cells increased depending on the noise effect (Fig. S1*B*-*E*), and resultant somites showed higher variance than those obtained in the presence of cell-cell interaction (Fig. S1*F*). Therefore, cell-cell interactions may play an important role in filtering ERK activity fluctuations caused by noise and in the robustness of ERK-mediated somitogenesis.

### Role of the Segmentation Clock

Finally, we removed the clock from the model (Fig. 5*A*). The ERK activity border alternated continuous movements and intermittent stops, at which somites were formed (Fig. 5*B* and *C*). Thus, the model predicted that stepwise ERK activity regression still occurred even without the clock, albeit with highly variable shift periodicity (Fig. 5*D*), leading to irregularly sized somite formation (110.4 ± 95.1 µm; C.V. = 0.86). Upon removal of cell-cell interactions in the absence of the clock, stepwise ERK activity regression was not observed (Fig. 5*E*), indicating that the high timing variations of the stepwise shift were caused by noise amplification through cell-cell interactions.

**Fig. 5.**
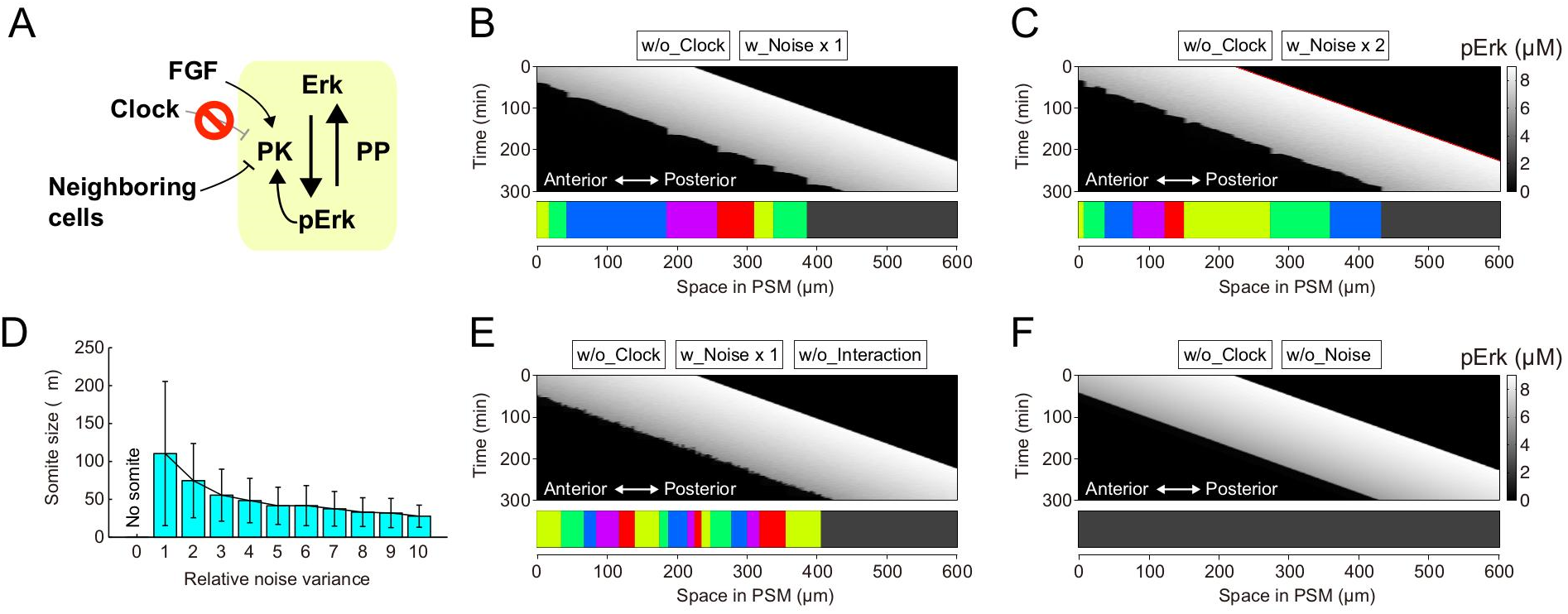
Stepwise ERK activity regression without the clock. (*A*) Simulation settings. Simulations were performed upon clock removal (i.e., *d* = 0 in Eq. 3). (*B, C)* Representative simulation results of ERK activity (upper panel) and resultant somites (lower panel) minus the segmentation clock with noise ×1 (B) and ×2 (C). (*D*) Noise effect on somite size without the clock. Formations of 200 somites were simulated by changing the noise variance from 0 to 10 times. Data represent the means and standard deviations. (*E, F*) Representative simulation results of Erk activity (upper panels) and the resultant somites (lower panels) without both clock and cell-cell interactions (*E*) and without both clock and noise (*F*). Neither stepwise Erk activity regression nor regular somite patterning occurred under these conditions.

The segmental defects in our model were reminiscent of those observed in clock-deficient embryos such as *her1* and *her7* double morphants and/or mutants (Fig. 1*C*) (10, 18, 19). However, these segmental defects could not be observed when we removed both the clock and the noise from our model; the ERK activity border continuously moved toward the posterior PSM and no somites formed (Fig. 5*F*), suggesting that intrinsic noise is utilized in ERK-mediated somitogenesis in living embryos. Based on these model predictions, we concluded that the clock has two distinct roles in somitogenesis: temporal regulation of stepwise ERK regression, and improvement of ERK-mediated somitogenesis developmental reproducibility.

### Noise-Resistant Somite Reproducibility by Clock

We investigated how somite reproducibility is generated against the noise. In the absence of cell-cell interactions and the clock, individual cell ERK activity levels differed owing to the noise (Fig. 5*E*). Thus, ERK activity up-down transitional timing inevitably differed among cells, leading to stepwise ERK activity regression failure. Adding cell-cell interactions locally coupled ERK activity among the neighboring cells, resulting reduced ERK activity homogeneity among neighboring cells (Fig. 5*B*). Because ERK activity regression timing differed owing to noise effects, the resultant somite size was highly variable (Fig. 5*D*). Therefore, our mathematical analyses suggest that cell-cell interactions contribute to filtering ERK activity homogeneities among neighboring cells, and that the filtering property adversely converts the noise effect within individual cells into ERK activity spatial heterogeneity (see Fig. 6*A* and Discussion for details).

**Fig. 6.**
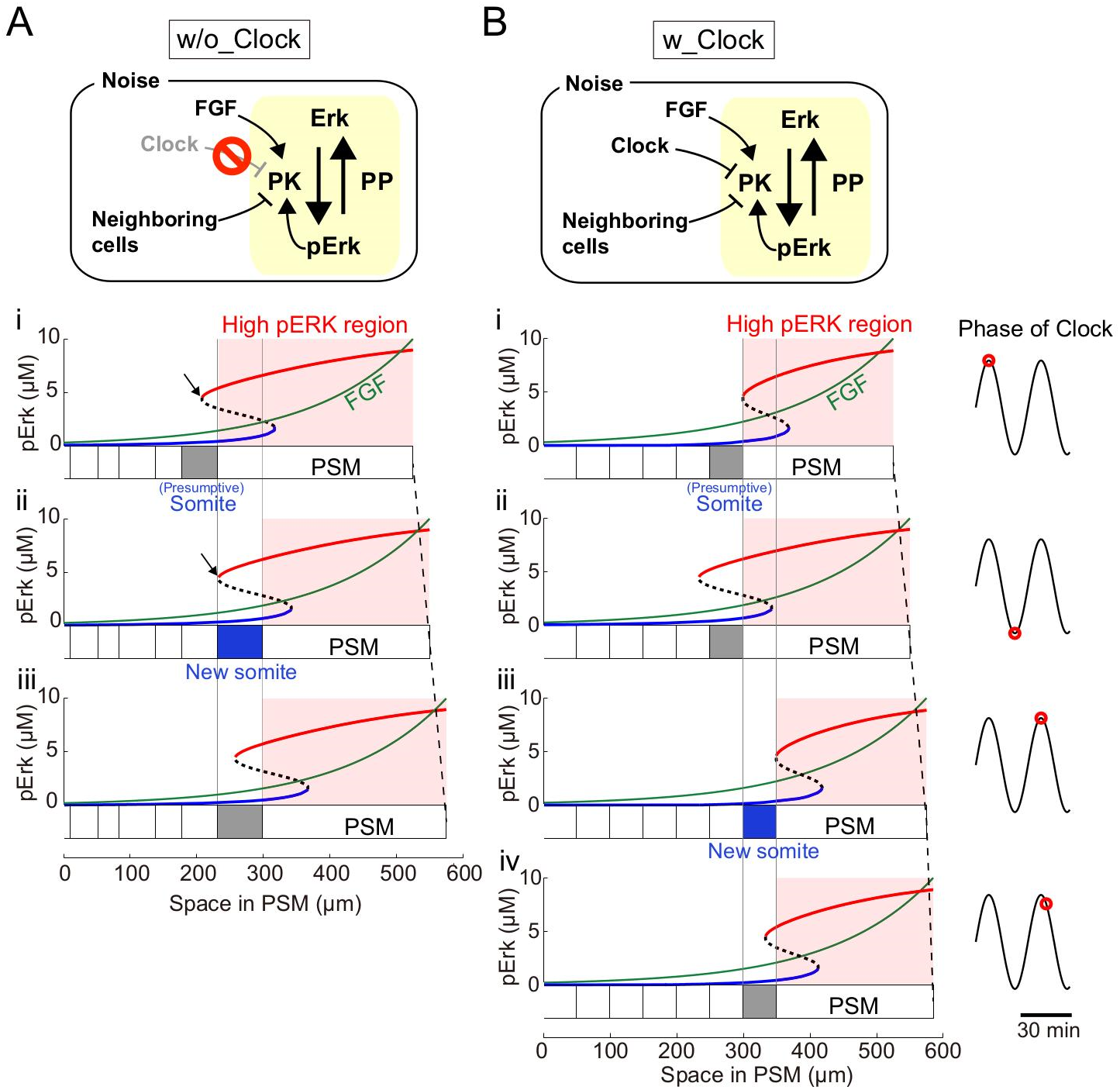
Mechanism of noise-induced irregular somite formation and clock-regulated somite reproducibility. Schematic representation of simulation conditions (upper panels). (*A*) Spontaneous irregular somite formation without the segmentation clock. (*i*) After a group of PSM cells forms the presumptive somite, the ERK activity bifurcation point localizes to the anterior presumptive somite (black arrow). (*ii*) Depending on FGF gradient posterior movement, the bifurcation point reaches the posterior presumptive somite border. Then, ERK activity rapidly transitions from the up to down state in undifferentiated cells, which enter the intermediate state. Cell-cell interaction propagates the ERK activity up-down state transition toward the posterior PSM in a chain-like manner (blue area), stopping at an irregular distance owing to the inevitable stochasticity. (*iii*) Process indicated in (*i*) is repeated after (*ii*). (*B*) Precise somite formation induced by the segmentation clock. (*i*) At peak clock phase, the clock-dependent ERK activity inhibition also peaks, drastically shifting the bifurcation diagram toward the posterior end [compare with (*A-i*)] and forming a new presumptive somite. (*ii*) At the mid-way clock phase point, the bifurcation diagram reverts to the anterior end [compare with (*A-ii*)], because the segmentation clock cannot inhibit ERK activity therein. (*iii*) Upon clock phase return, the bifurcation diagram shape gradually shifts toward the posterior end [compare with (*B-i*)]. During this phase, a PSM cell group transitions from the up to down ERK activity state, resulting from temporal oscillation and spatial bifurcation diagram shift, due mainly to the segmentation clock and PSM growth, respectively. Also, the up-to-down ERK activity transition tend to propagate toward the posterior end owing to the inevitable stochasticity, which may lead to somite irregularities as with segmentation clock loss [(*A-ii*) and Fig. 1*C*]. (*iv*) Immediately thereafter, the propagation terminates at a fixed distance, because the bifurcation diagram starts to shift toward the anterior end [compare with (*B-iii*)].

Adding the clock into our model reduced ERK activity spatial heterogeneity, which is caused by the noise and cell-cell interactions, leading to reproducible constant-sized somite formation (Fig. 2*D*). To understand its mechanism, segmentation clock effects on the PSM ERK activity bifurcation diagram were investigated. Because of the clock, the ERK activity bifurcation diagram cyclically shifted back-and-forth in the PSM (see Fig. 6*B* and Discussion for details). After each clock cycle, the bifurcation diagram shifted to the posterior end owing to PSM elongation. Thus, the ERK activity up-down transition was precisely restricted to a specific time and within specific space intervals. Therefore, these results suggest that the segmentation clock both provides temporal information on somite formation as in the classical CW model and reduces ERK activity spatial heterogeneity against the noise, representing a previously unidentified role of the segmentation clock.

## Discussion

The mechanisms underlying the precision and reproducibility of developmental events within embryos in multicellular organisms despite cell-to-cell variations in mRNA and protein levels caused by stochastic gene expression (21) remain unclear. Through computational modeling, we clarified the mechanism by which high reproducibility among somites is achieved. We also proposed a novel concept wherein temporal regulation of the global signal; i.e., the segmentation clock, reduces the noise effect to enhance developmental reproducibility.

### Model Prediction

The CW model predicted that clock removal impeded somite individualization, as seen in simulation without clock and noise (Fig. 5*F*). This contradicted clock gene loss-of-function phenotypes wherein irregularly-sized somites are still formed. In contrast, our model has an ability to generate such somites, predicting that stepwise ERK activity regression still occurred at irregular timing with irregular distances in clock-deficient embryos (Fig. 5*B*-*C*). This may explain why stepwise ERK activity regression could not be observed in clock-deficient embryos in our previous static analysis based on collected ERK activity snapshots from different embryos (18). Therefore, live ERK activity imaging within an embryo is needed to validate our prediction.

### Comparison with Previous Models

The CW model, originally proposed as a conceptual model (8), was revised upon clock and wavefront molecular bases discovery (22, 23). Several theoretical models have also been proposed to explain the different aspects of somitogenesis, including the mechanism of PSM cell oscillatory gene expression (24, 25), clock synchronization among cells (26–29), and PSM spatial oscillatory patterning (30). Although two models addressed the noise effect on clock synchronization (26, 29), none addressed that on somite commitment. Thus, our model is the first to satisfactorily explain how somite reproducibility is maintained subject to noise.

### Validity of Model Components

Our model exhibits biological relevance in terms of the intrinsic noise, ERK signaling bistability, cell-cell interactions, and the segmentation clock.

#### Intrinsic Noise

In our model, added noise perturbed ERK activity. Physically, chemical reactions happen stochastically, subjecting molecular concentrations to random fluctuations when molecules are present in low copy numbers (1–3). Synthetic biological studies on bacteria and yeast demonstrate the inevitability of reaction noise in gene expression regulation (1, 3). Although it is difficult to identify the noise source and manipulate its intensity in the developing embryo, we considered that it originates from intracellular reaction stochastic properties within single cells.

#### ERK Signaling Bistability

In a concept of canalization proposed by Waddington (31), the cell state follows a downward slope to reach a stable point associated with cell fate transition. In a current interpretation, cell fate transitions are defined by stable gene expression patterns. Accordingly, previous somitogenesis models incorporated two stable states (i.e., bistability, which associates with cell fate transition during somite individualization) (8, 22). Because we previously observed up and down PSM ERK activity states (Fig. 1*G*), and as *Xenopus* oocyte ERK activity exhibits bistability, our model incorporated ERK signaling bistability regulated by positive feedback (32).

#### Cell-Cell Interaction

Somite formation is a type of collective cell fate transition in a population of cells, suggesting that cell-cell interactions are involved in somite formation. It has been reported that the spontaneous formation of somite-like structures in the absence of the clock requires cell-cell interactions (20); moreover, PSM cell ERK activity levels are similar to those of adjacent PSM cells (18). Thus, our model assumed cell-cell interactions, such that cells in the ERK activity down state down-regulate neighboring cell ERK activity (Fig. 2*A*).

#### Segmentation Clock

No direct evidence supports that clock genes (*her1* and *her7*) temporally modulate zebrafish ERK activity. However, negative ERK regulator (e.g., *sprouty* and *dusp*) mRNA expression oscillates in the mouse and chick (9). Thus, we considered that the segmentation clock temporally regulates PSM ERK activity. In the model, we tentatively assumed a negative clock effect on ERK activity (Fig. 2*A*). However, the model reproduced stepwise ERK activity regression regardless of clock effect direction (Fig. 2*D* and Fig. S2), suggesting the relevance of clock periodicity to signaling but not effect direction.

### Clock-Independent and -Dependent Somite Formation Mechanism

Finally, we proposed mechanisms for clock-independent irregular- and clock-dependent regular somite formation from the point of view of a dynamic system subject to noise. Our model predicted irregular ERK activity border shifting in clock-deficient embryos (Fig. 5*B* and *C*). Here, we depicted a bifurcation diagram in space within the growing PSM (Fig. 6A). A newly generated cell at the posterior PSM end is exposed to a high FGF gradient, which activates ERK (Fig. 6*A*, also see Fig. 2*B*). The cell with the up ERK activity state then aligns toward the anterior PSM alongside tail elongation, leading to gradual FGF gradient decrease. Owing to ERK signaling bistability, ERK remains activated; when the cell reaches a specific point in the bifurcation diagram (black arrow in Fig. 6*A**-ii*), ERK suddenly inactivates. This cell then starts to interact with and induce neighboring cells to undergo up-down ERK activity transition, which propagates toward the posterior PSM. Owing to the inevitable stochasticity, this propagation progresses stochastically and terminates if the up-down ERK activity state transition does not happen within a specific time interval (Fig. 6*A**-ii*, *blue region*). Thus, the ERK activity border shifts an irregular distance, eventually leading to irregularly sized somite formation.

Given the clock-independent mechanism of irregular somite formation described above, we investigated the segmentation clock role in reducing the noise effect on somite reproducibility. With clock present, the ERK activity bifurcation diagram cyclically changed depending on the clock phase (Fig. 6*B*). This may explain how the regular somite pattern is generated against the inevitable stochasticity. Within a specific time interval (before peak clock oscillation), ERK is inactivated in an anterior PSM cell cohort (Fig. 6*B**-iii*). At this time, the ERK activity up-down transition starts to propagate toward the posterior PSM, as seen upon segmentation clock loss (Fig. 6*A*). Immediately thereafter (after peak clock oscillation), however, the bifurcation diagram shifts toward the anterior end, leading to stochastic propagation termination at a fixed distance (Fig. 6*B**-iv*). Accordingly, uniformly sized somites are periodically generated.

## Methods

### Computational Model for ERK-Mediated Somite Formation

A computational model of somite segmentation that incorporates the FGF gradient, ERK signaling, segmentation clock, cell-cell interaction, and reaction noise was developed.

In the PSM, FGF is produced at the posterior end and degraded over time. Depending on tail elongation, PSM cells are added to the posterior end so that they gradually move toward the anterior end (Fig. 2*B*) (6). Thus, the FGF concentration at position *x* (the origin is fixed at a specific anterior point) is represented by

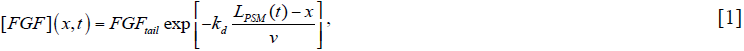

where *FGF_tail_*, *k_d_*, and *v* denote the tail peak concentration, FGF degradation rate, and tail elongation speed, respectively. PSM elongation is represented by *L*_*PSM*_(*t*) = *L_o_* + *vt*, where *L_o_* is the initial PSM length. The equation (*L_PSM_*(*t*) – *x*)/*v* represents spent time of the cell at position *x* from birth at the posterior PSM end. A one-dimensional PSM cell array is assumed in the model.

ERK is activated and inhibited by protein kinase (PK) and protein phosphatase (PP), respectively. pERK dynamics in each cell are described by

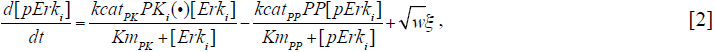

where *i* indicates the cell index; *kcat_l_* and *Km_l_* (*l* ∈ {*PK*, *PP*}) are catalytic reaction rates and Michaelis-Menten constants, respectively; *PK_i_*(•) is the active protein kinase concentration described by equation [3] and *PP* is active protein phosphatase concentration; ξ_i_ and *w* indicate the white noise 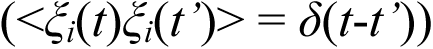 and noise variance, respectively. The total ERK amount is assumed to be constant as described by *ERK_tot_* = [*ERK_i_*] + [*pERK_i_*].

PK is regulated by positive feedback from pERK, the FGF gradient, the segmentation clock, and cell-cell interaction by

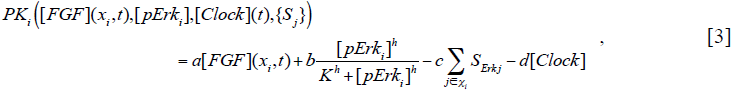

where *a*, *b*, *c*, and *d* indicate positive constants; *K* and *h* indicate [*pERK*] concentration required for the half-maximal activation of positive feedback and Hill coefficient, respectively; *χ_i_* indicates the set of neighboring cells of cell *i*; and *S_j_* is the binary value indicating the intermediate state of the neighboring cell *j*. For intermediate state cells (i.e., with a down ERK activity state), *S_j_* = 1; otherwise (i.e., undifferentiated or differentiated states), *S_j_* = 0. [*Clock*] indicates clock gene cyclic expression, assumed to be spatially uniform over the PSM by

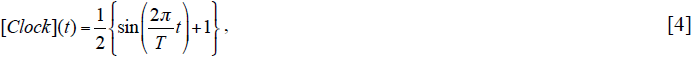

where *T* is the oscillation period.

The parameters used in the model were as follows: *FGF_tail_* = 4 (µM), 1/*k_d_* = 1.5 (h), *v* = 100 (µm/h), *ERK_tot_* = 10 (µM), *kcat_PK_* = 10 (s^−1^), *kcat_PP_* = 10 (s^−1^), *Km_PK_* = 5 (µM), *Km_PP_* = 0.7 (µM), *PP* = 0.5 (µM), *K* = 3 (µM), *h* = 4, *w* = 0.3 (µM/s), *T* = 30 (min), *a* = 0.7, *b* = 0.5, *c*=0.05, and *d* = 0.2. For simulation, the pERK threshold between undifferentiated and intermediate states was set to 3 (µM). The time interval required for the intermediate to differentiated state transition was set at 200 (s). These values were selected based on three criteria as follows: (1) FGF gradient formed in the PSM as shown in Fig. 2*B* (for *FGF_tail_*, *k_d_*, and *v*), (2) PSM pERK exhibited a bistable bifurcation diagram depending on the FGF gradient as shown in Fig. 1*B* (for *ERK_tot_*, *kcat_PK_*, *kcat_PP_*, *Km_PK_*, *Km_PP_*, *PP*, *K*, *h*, *a*, *b*), and (3) segmentation clock cyclically changed the bifurcation diagram as shown in Fig. 5*B* (from *i* to *iii*). Under conditions that satisfied these criteria, the basic behaviors shown in Fig. 2*D* and Fig. 5*B* were robustly reproduced.

## Acknowledgments

We are grateful to Michiyuki Matsuda for offering advice, helpful discussions, and critically reading the paper. This work was partially supported by Grants-in-Aid for Scientific Research from the Ministry of Education, Culture, Sports, Science and Technology (MEXT), Japan (to H.N., R.A., S.I., Y.B., and T.M.), the Takeda Science Foundation (to T.M.), and the Platform Project for Supporting Drug Discovery and Life Science Research (Platform for Dynamic Approaches to Living System) from the Japan Agency for Medical Research and Development (AMED) (to S.I. and H.N.).

## Author contributions

H.N. and T.M. conceived the project. H.N. developed the model and performed the computational simulation. H.N. and T.M. wrote the draft, and H.N., T.M., R.A., S.I., and Y.B. prepared the final version of the manuscript.

## Supplemental Figures

**Fig. S1.**
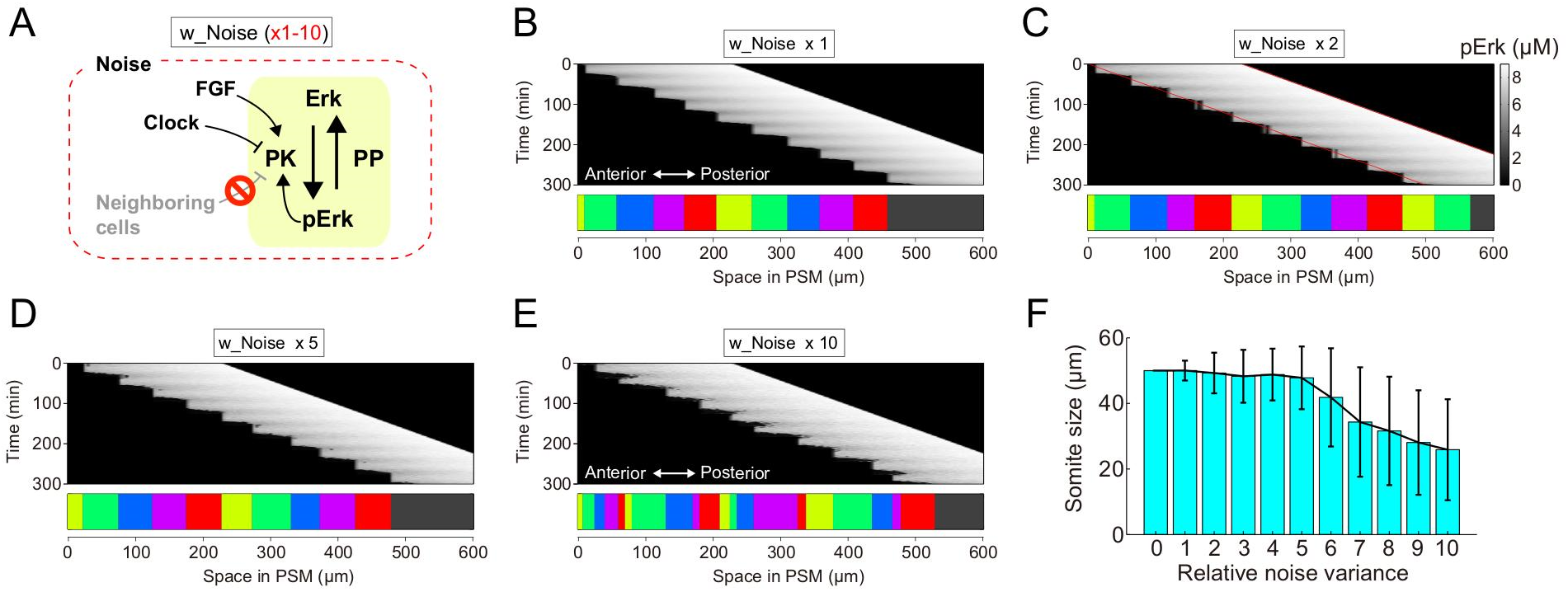
Erk-mediated somitogenesis without cell-cell interactions. (*A*) Simulation settings. Simulations were performed for cell-cell interaction exclusion (i.e., *c*=0 in Eq. 3). (*B-E*) Representative simulation results of Erk activity (upper panel) and the resultant somites (lower panel) when the noise variance was set to standard conditions (*B*) and increased 2 (*C*), 5 (*D*), or 10 (*E*) fold. (*F*) Effect of noise on somite size. Formations of 200 somites were simulated by changing the noise variance from 0 to 10 fold. The data represent the means and standard deviations.

**Fig. S2.**
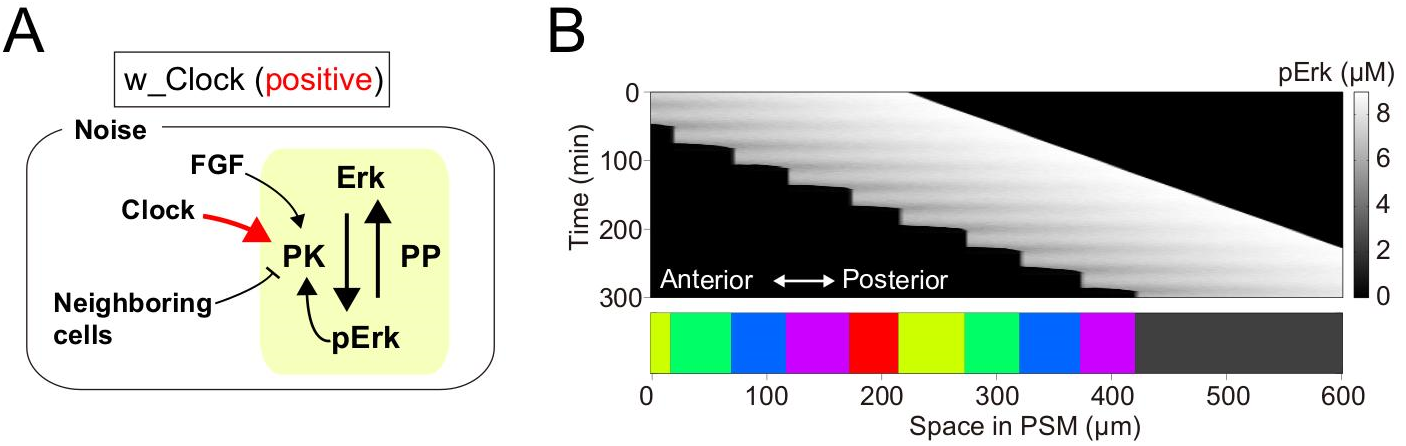
The case of positive clock effect on the ERK activity. (*A*) Simulation settings. Simulations were performed for a positive clock effect (i.e., *d* = −0.2 in Eq. 3). (*B*) Representative simulation result of Erk activity (middle panel) and resultant somites (lower panel). Note that somite formation occurs normally, even when Erk is positively regulated by the clock.

